# Droplet microfluidics for time-resolved serial crystallography

**DOI:** 10.1101/2024.01.12.575388

**Authors:** Jack Stubbs, Theo Hornsey, Niall Hanrahan, Luis Blay Esteban, Rachel Bolton, Martin Malý, Shibom Basu, Julien Orlans, Daniele de Sanctis, Jung-uk Shim, Patrick D. Shaw Stewart, Allen M. Orville, Ivo Tews, Jonathan West

## Abstract

Serial crystallography requires large numbers of microcrystals and robust strategies to rapidly apply substrates to initiate reactions in time-resolved studies. Here we report the use of droplet miniaturisation for the controlled production of uniform crystals, providing an avenue for controlled diffusion and synchronous reaction initiation. The approach was evaluated using two enzymatic systems, yielding 3-µm lysozyme crystals and 2-µm crystals of Pdx1, an Arabidopsis enzyme involved in vitamin B6 biosynthesis. A seeding strategy was used to overcome the improbability of Pdx1 nucleation occurring with diminishing droplet volumes. Convection within droplets was exploited for rapid crystal mixing with ligands. Mixing times of <2 milliseconds were achieved. Droplet microfluidics for crystal size engineering and rapid micromixing can be used to advance time-resolved serial crystallography.

## 1. Introduction

Modern crystallography incorporates diffraction data collection at room temperature, providing a means to emulate physiological conditions whilst also observing the dynamic nature of proteins (Orville, 2020; Fraser *et al*., 2011, Fischer, 2021). Challenges posed by elevated radiation damage (Holton, 2009; Garman, 2010; Garman & Weik, 2023), can be overcome by the collection of multiple datasets or the application of serial methods. No longer can optimal crystals be hand-picked, but instead large numbers of uniform microcrystals must be prepared. Advancements in instrumentation, including high flux synchrotron sources and extreme brilliance X-ray free electron lasers (XFELs) (Chapman et al., 2011; Chapman *et al*., 2019; Barends *et al*., 2022), coupled with developments in automation, data processing, detector technologies (Forster *et al*., 2019) and sample delivery, will ensure time-resolved experiments using serial methods will become routine in the near future.

Crystal size is a critical parameter for many reasons (Shoeman *et al*., 2022). Crystal size should be tuned to the synchrotron or XFEL beam size (∼1–20 µm (Evans *et al*., 2011)) for improved signal-to-noise ratio in the X-ray diffraction pattern and efficient use of the protein sample. For time-resolved studies, the key advantages of small crystals are short substrate transport paths into the crystal lattice for rapid mixing or short light paths for full penetration of exciting light. For illustration, substrate diffusion into the centre of a 2-mm crystal (*i*.*e* 1 mm travel) is dependent upon several factors, including ligand diffusion coefficient, initial concentration, charge, mother liquor viscosity, and crystal lattice packing, with time scales ranging from 400 ms for O_2_ (32 Da) to 3.5 ms for larger ligands (*e*.*g*. ceftriaxone, 554 Da) (Schmidt, 2013). Crystal uniformity is critically important, especially for synchronised reaction triggering, but also to avoid large crystals clogging capillaries used in many sample delivery systems. Ideal results will derive from monodisperse microcrystal slurries, robust sample delivery methods, and reaction initiation strategies that exploit the particular X-ray source characteristics and limit sample consumption.

Preparing large numbers of uniformly small crystals is an on-going challenge for the field. While microcrystal showers are often the first hit in sparse matrix vapour diffusion screens, they typically need to be scaled-up by batch methods to produce the volumes required for serial crystallography experiments; this may reach millilitre volumes for time-course experiments with multiple time-point datasets (Beale *et al*., 2019; Stohrer *et al*., 2021; Beale & Marsh, 2021; Shoeman *et al*., 2022).

Crystal formation typically comprises a nucleation phase, followed by a growth phase. In some crystallisation conditions nucleation occurs rapidly, and as crystals grow they deplete protein in solution, and thereby prevent further nucleation. However, this results in a variation of crystal sizes; for example, if two crystal nuclei from in close proximity the competition for protein will result in a pair of smaller than average crystals. A popular strategy is, therefore, to fragment crystals to make a seed stock that can be used to control crystal growth (Stura & Wilson, 1990; D’Arcy *et al*., 2007; Shaw Stewart *et al*., 2011; Shoeman *et al*., 2022). By increasing the number of seeds, crystal size can be reduced.

Microfluidics has attracted significant attention for crystallography as it can precisely control reaction environments (Li & Ismagilov, 2010; Puigmartí-Luis, 2014; Shi *et al*., 2017; Sui & Perry, 2017). Initial efforts involved nanoliter environments enabling counter-diffusion for exploring phase diagrams (Hansen *et al*., 2002; Zheng *et al*., 2004; Li & Ismagilov, 2010), or dialysis to decouple and optimise nucleation separately from growth (Shim *et al*., 2007, Shim *et al*., 2007; Selimovic *et al*., 2009). Droplet microfluidic formats then allowed better control of crystal formation by negative feedback through protein depletion during crystal growth, (Dombrowski *et al*., 2007; Heymann *et al*., 2014) defining crystal size by available protein, *i*.*e*. droplet volume. Droplet microfluidic crystallisations have been demonstrated for lysozyme, glucose isomerase, trypsin, concanavilin A, D1D2 splicesomal snRNP particle (Heymann *et al*., 2014; Akella *et al*., 2014), sugar hydrolase and sialate O-acetylesterase (Babnigg *et al*., 2022). Importantly, microfluidic droplets are highly monodisperse, which allows the protein supply to be exactly metered to achieve crystal uniformity. Studies to date have optimised droplet size to achieve single crystal occupancy for the formation of large crystals suitable for obtaining synchrotron diffraction data *in situ*.

For time-resolved experiments there is also the challenge of rapidly triggering reactions with substrates and ligands (Echelmeier *et al*., 2019). Mix-and-inject methods first emerged (Weierstall *et al*., 2012; Calvey *et al*., 2016; Olmos *et al*., 2018; Ishigami *et al*., 2019; Dasgupta *et al*., 2019; Pandey *et al*., 2021; Murakawa *et al*., 2022) that involve coaxial flows with a core crystal stream. Hydrodynamic focussing results in stream thinning to provide short paths for the diffusion of substrate molecules into the crystal prior to high-velocity injection into the beam using a gas dynamic virtual nozzle (GDVN) (DePonte *et al*., 2008). As an efficient, high hit rate alternative, piezoelectric or acoustic drop-on-demand methods are gaining popularity for the delivery of substrate droplets onto crystals presented on fixed targets (Mehrabi *et al*., 2019) or tape drives (Roessler *et al*., 2016; Fuller *et al*., 2017, Butryn *et al*., 2021). Here, picolitre substrate volumes are dispensed onto individual crystals or crystals contained in nanolitre droplets (both within a humidified environment). Mixing initially occurs by impact-induced convection, followed by diffusion, then the registration of the crystal into the beam after a defined time delay (2 ms and upwards). These sample delivery methods and their considerations are captured in recent reviews (Schulz *et al*., 2022; Barends *et al*., 2022).

In this contribution, we explore droplet scaling from nanoliter volumes down to sub-picoliter volumes and demonstrate the ability to engineer crystal size and uniformity. Using *Arabidopsis thaliana* Pdx1, an enzyme involved in vitamin B6 biosynthesis (Rodrigues *et al*., 2017, 2022), and lysozyme, we demonstrate crystal scaling to suitable dimensions and numbers for time-resolved serial crystallography. We show that with diminishing volumes, nucleation becomes improbable, but this can be countered by seeding. We go on to exploit droplets as convective environments for rapid micromixing, achieving mixing within 2 milliseconds to support future strategies for understanding structural dynamics with high temporal resolution.

## 2. Material and Methods

### 2.1 Protein expression, purification and crystallisation

#### 2.1.1 Lysozyme

Lysozyme (chicken egg white, Melford) was batch crystallised using a ratio of 1 part 20 mg/mL lysozyme in 20 mM sodium acetate, pH 4.6 to 4 parts of mother liquor; 6% PEG 6000 (w/v) in 3.4 M NaCl and 1 M sodium acetate, pH 3.0; (adapted from previous conditions (Martin-Garcia *et al*., 2017)). The mixture was vortexed for 5 seconds and left to crystallise for 1 hour at room temperature.

#### 2.1.2 Trypsin

Trypsin (bovine pancreas, type I, Merck) needles were crystallised using seeded vapour diffusion conditions as previously described (Heymann *et al*., 2014). Seed stocks were prepared by pooling crystals from many vapour diffusion drops, dilution in mother liquor (11–14% PEG 4000 (w/v), 15% ethylene glycol, 200 mM SiSO_4_, 100 mM MES, pH 6.5) and vortexing with a Hampton Seed Bead for 180 seconds by alternating between 30 seconds of vortexing and 30 seconds on ice followed by storage at -20°C. Seeded-batch trypsin crystallisation involved 1 part of 65 mg/mL trypsin in 3 mM CaCl_2_ with benzamidine, 1 part mother liquor and 1 part seed prepared in mother liquor. The mixture was vortexed for 5 seconds and incubated at room temperature overnight.

#### 2.1.2 Pdx1

Plasmid encoding wild-type Pdx1.3 (UniProt ID: Q8L940; EC:4.3.3.6; Rodrigues *et al*., 2017) was transformed into BL21 (DE3) competent *E*.*coli* cells, grown to OD_600_ 0.6 at 37°C. After induction with 25% (w/v) lactose and growth for a further 16 hours at 30°C, cells were harvested by centrifugation. Cells were resuspended in lysis buffer (50 mM Tris pH 7.5, 500 mM sodium chloride, 10 mM imidazole, 2% (v/v) glycerol) and sonicated on ice. The lysate was ultracentrifuged at 140,000 × g at 4°C for 1 hr, filtered and immobilized on a metal ion affinity chromatography HisTrap HP column (GE Healthcare). Pdx1 was washed and eluted with lysis buffer, containing 50 mM and 500 mM imidazole respectively, as well as 5% (v/v) glycerol. The eluted protein was buffer exchanged into gel filtration buffer (20 mM Tris pH 8.0, 200 mM KCl), before centrifugal concentration with a 30 kDa cut-off (Vivaspin 20, Sartorius). Crystals for preparing seeds were produced by combining (1:1) ∼12 mg/mL Pdx1 with mother liquor (600 mM sodium citrate, 100 mM HEPES pH 7) as 10 L vapour diffusion drops in 24-well XRL plates (Molecular Dimensions). Crystals for seed stocks grew overnight and varied in size from 10–50 μm in length. Vortexing with a Hampton Seed Bead was carried out for a total of 180 seconds by alternating between 30 seconds of vortexing and 30 seconds on ice followed by storage at -20°C. Batch crystallisation involved a 1:1:1 mixture of 12 mg/mL Pdx1, seed (10^5^–10^7^/mL) and mother liquor. In droplets a 2:1 mixture of seeds (10^7^/mL) diluted in mother liquor with 12 mg/mL Pdx1 was used.

### 2.2 Droplet Microfluidics

#### 2.2.1 Device Fabrication

Microfluidic devices (Whitesides, 2006) were replicated by standard soft lithography (Whitesides, 2001) using SU-8 wafers for the moulding of poly(dimethylsiloxane) (PDMS, Sylgard 184) devices with curing at 60°C for 2 hours. A range of different droplet microfluidic devices were used for the generation of nanolitre to femtolitre droplets. Droplet generation junction dimensions, flow rates and droplet characteristics for the different protein systems are documented in Table S1-S3. Tubing ports were introduced using 1-mm-diameter Miltex biopsy punches (Williams Medical Supplies Ltd). Devices were bonded to glass microscope slides using a 30 s oxygen plasma treatment (Femto, Diener Electronic) followed by channel surface functionalization using 1% (v/v) trichloro(1*H*,1*H*,2*H*,2*H*-perfluorooctyl) silane (Merck) in HFE-7500™ (3M™ Novec™).

#### 2.2.2 Experimental Setup

The experimental set-up for droplet generation is shown in Figure S1. The process involved the preparation of syringes containing, protein, mother liquor and fluorinated oil (QX200™, BioRad) which acts as the carrier phase. Pdx1 and trypsin I droplet preparations required the use of seeds within the mother liquor. Syringes were interfaced with 25 G needles for connecting to the microfluidic ports via polythene tubing, (ID 0.38 mm; OD 1.09 mm, Smiths Medical). Syringe pumps (Fusion 100, Chemyx) were used to deliver reagents for microfluidic droplet generation. Droplet generation was monitored using a Phantom Miro310 (Ametek Vision Research) high-speed camera mounted on an inverted microscope (CKX41, Olympus). Droplets were collected in microcentrifuge tubes and stored at room temperature for 2–3 days with a mineral oil overlay to prevent coalescence. Droplet dimensions and crystal occupancy were measured using a supervised ImageJ (NIH) process. Lambda (⍰) is used to denote the average number of crystals per droplet.

#### 2.2.3 Crystal retrieval and analysis

Crystals were retrieved from droplets by a procedure called *breaking the emulsion*. First, the QX200™ oil is removed, then a 10-fold volume (relative to emulsion volume) of mother liquor is added. Next, a volume of 1*H*,1*H*,2*H,2*H-perfluoro-1-octanol (PFO, Merck) is added to the emulsion with gentle pipetting used to break the emulsion. The PFO exchanges with the commercial surfactant surrounding the droplets, allowing the aqueous compartments of droplets to contact each other and coalesce. Finally, the single aqueous volume containing the crystals is removed for analysis by mounting on a coverslip for oil immersion imaging with a 60x/1.4NA objective (Olympus). Crystal dimensions were measured using a custom MATLAB script (https://github.com/luiblaes/Crystallography) and manually validated. Crystal samples were taken to the European Synchrotron Radiation Facility (ESRF) for serial synchrotron measurements.

### 2.3 Serial synchrotron crystallography (SSX)

Ground state structures were obtained by SSX. Lysozyme and Pdx1 crystals were concentrated by settling and applied to sheet-on-sheet (SOS) chips (Doak *et al*., 2018). This involved removal of excess liquid and sandwiching 3–5 µL of the crystal slurry between two Mylar^®^ films and sealing inside a metal mount. A total of 81,800 images were collected per foil. SSX data for lysozyme and Pdx1 crystals grown in batch and within microfluidic droplets were collected on the new ID29 beamline at the European Synchrotron Radiation Facility (ESRF, France) using a 2×4 m (VxH) beam of 11.56 keV X-rays, with a 90 s pulse and 231.25 Hz repetition rate, and a 20 μm step movement between images. A JUNGFRAU 4M detector (Mozzanica *et al*., 2018) with a sample-to-detector distance of 175 mm (1.8 Å in the corner) was used to collect diffraction patterns. Full data collection and processing details are documented in the Supplementary Information.

### 2.4 Mixing in droplets and image analysis

Mixing of lysozyme crystals (7×2 mm; grown by batch crystallisation) with 25 mM sulfanilic acid azochromotrop (SAA, Merck, l_max_ 505–510 nm), a highly absorbing red dye, was investigated using 30×40 ⍰m droplet generation junctions with an oil:aqueous flow ratio of 2:1. The crystal:dye flow ratio was modulated along with total flow rates ranging from 7.5 to 45 mL/min. To retain crystals in suspension for ensuring continuous crystal delivery to the microfluidic device we used automated syringe rotation (Lane *et al*., 2019). In an alternative setup, a droplet generator producing SAA droplets was positioned upstream of an inlet for the introduction of pre-formed ∼70-⍰m-diameter droplets containing lysozyme crystals. The lysozyme and SAA droplets were synchronised for one-to-one interception, followed by surfactant exchange with PFO for droplet fusion and ensuing circulation-driven micromixing. This experiment involved 12.5 ⍰L/min 10% (v/v) QX200 in HFE7500, 4 ⍰L/min lysozyme, 5 ⍰L/min SAA dye and 4 ⍰L/min PFO flow rates. For both strategies, diffusive-convective mixing of the SAA dye was captured by hi-speed imaging (Phantom Miro310, Ametek Vision Research). Droplets were individually analysed to understand mixing with and without crystals. The coefficient of variation (CV) of the intensity of pixels defining each droplet was used as the mixing measure. The CV approaches zero as the dye is homogenised throughout the droplet. The time from stream combination to a 5% pixel intensity CV value was used to define the mixing time. Mixing analysis was automated using a MATLAB script with 15 single droplet kymographs used to derive mixing time statistics.

## 3. Results

### 3.1 Experimental Design

Droplet microfluidic designs incorporated aqueous inlets for protein, mother liquor, seed and another for the fluorinated oil with flow focussing used for droplet generation [Fig. 1(a)]. Droplet generation junction dimensions were used to scale droplet volumes from ∼200 pL to ∼1 pL to investigate conditions for controlling lysozyme and Pdx1 crystal size and uniformity. To understand the effects of droplet confinement, resulting crystals were compared with those grown under conventional batch conditions. Droplet microfluidics was then investigated as a means to rapidly mix crystals with substrates. Crystals were either encapsulated with substrate during droplet generation or crystal-containing droplets were fused with substrate-containing droplets.

**Figure 1.**
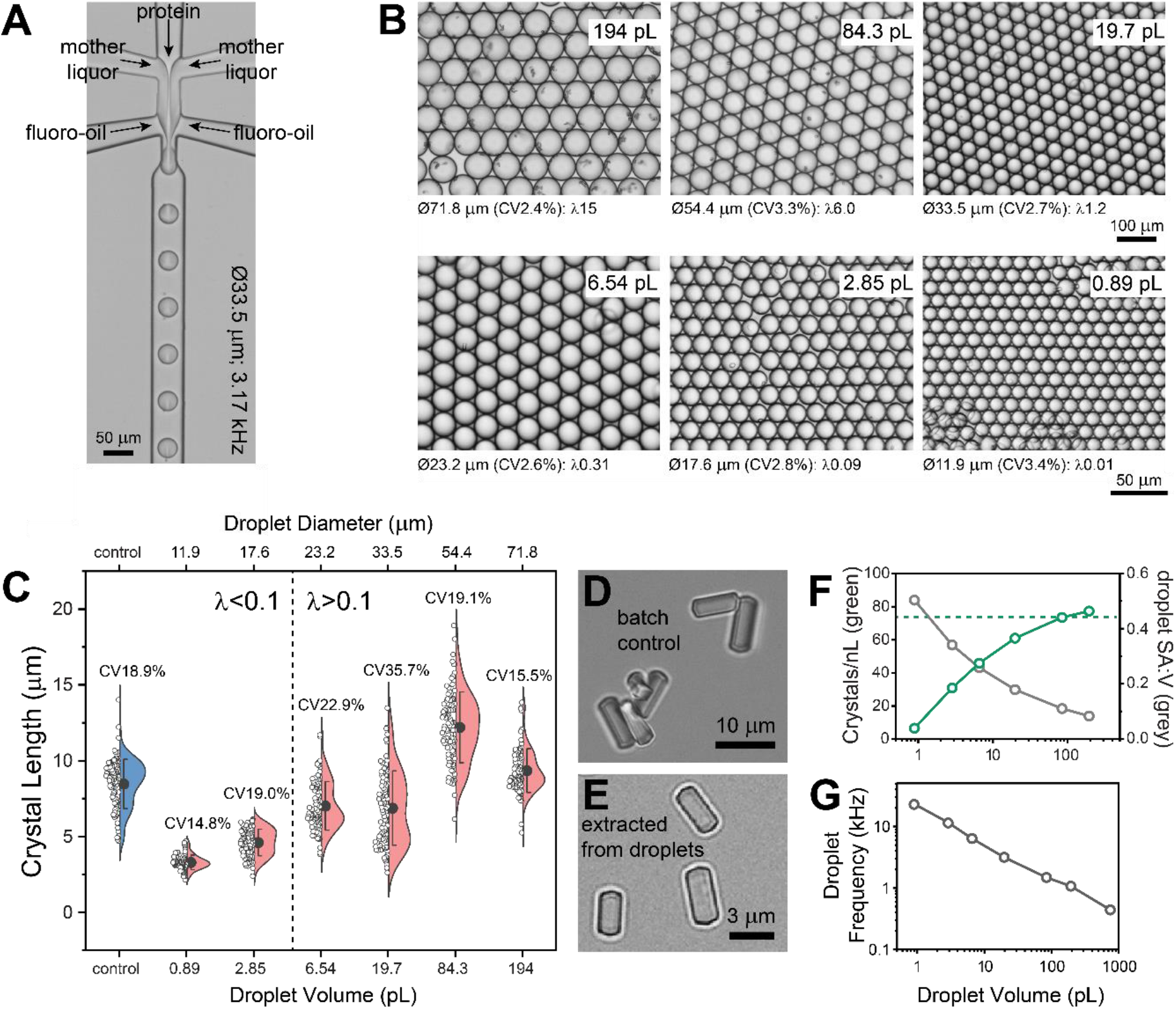
Lysozyme crystal size control by droplet volume scaling. (a) Protein crystallisation droplets generated at kHz frequencies by combining streams of lysozyme, mother liquor and fluorinated oil. (b) Using different devices and flow rates (see SI table 1), monodisperse droplets (CV<4%) can be produced with picolitre to femtolitre volumes. (c) Lysozyme crystals produced in batch conditions (control, blue) were on average 8-⍰m-long. The length of lysozyme crystals produced in droplets (salmon) correlates with droplet volume, with ∼3-⍰m-long crystals produced in the smallest 0.89 pL droplets. Crystal uniformity emerges when the average number of crystals per droplet (⍰) is ≤0.1. (d,e) Visual comparison of lysozyme crystals prepared in batch (control) and extracted from 0.89 pL droplets by breaking the emulsion. (f) Droplet volume miniaturisation is associated with reduced crystal density normalised to crystals/nL (green) which correlates with increasing surface area to volume (SA:V, grey) ratio. The batch crystal density value is denoted by the green dashed line. (g) Gains in droplet generation frequency scale with droplet volume reduction.

### 3.2 Lysozyme crystallisation in microfluidic droplets

Lysozyme is a well-known standard that undergoes extremely fast nucleation (Forsythe *et al*., 1999). Indeed, the nucleation rate in our batch crystallisation method is too fast to measure (Video S1), but a resultant crystal density of ∼80/nL was observed (∼80M/mL). The rapid growth of lysozyme crystals introduces negative feedback to prevent later nucleation events. This aids length uniformity, producing crystals with an 8 ⍰m average length and a coefficient of variation (CV) of ∼19% [Fig. 1(c)].

**Table 1.**
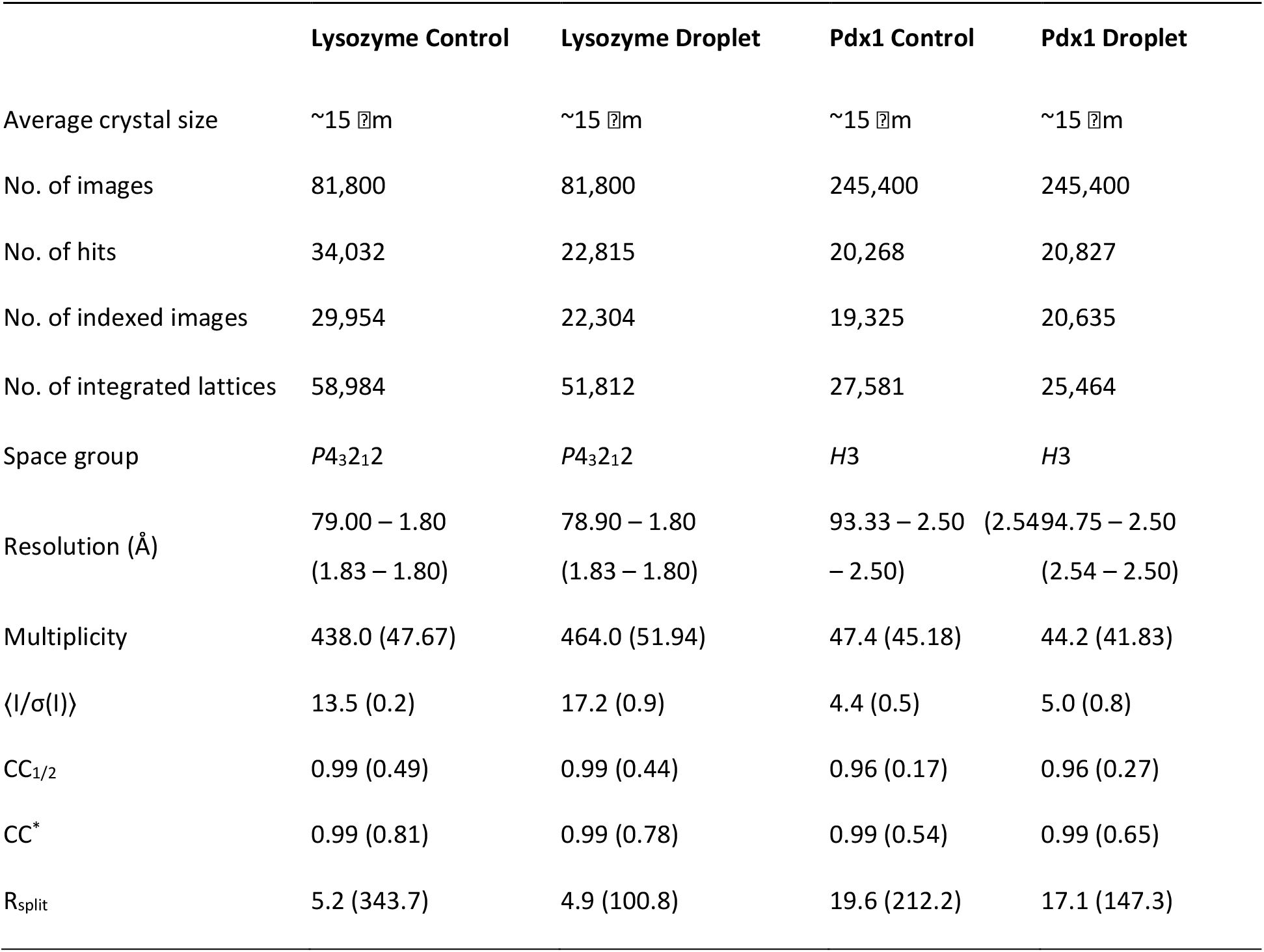
Data collection statistics for lysozyme and Pdx1 crystals grown in batch conditions and in droplets. Data collected at ESRF/ID29. Full data is reported in Table S4. Values in parentheses are for the outermost resolution shell.

Using batch crystallisation as a benchmark we then sought to understand the effects of volume scaling by droplet confinement. Droplet microfluidics produced monodisperse (CV<4%) droplets ranging in size from 754 to 0.89 pL and crystal sizes ranging from 20 mm to 2 mm [Figs. 1(a) and 1(b)]. In the largest droplets (754 pL (<u>Ø</u>113 mm)), crystals were too numerous to count, whereas smaller droplets showed a crystal occupancy ranging from an average of 15 crystals/droplet (⍰15) in 194 pL droplets to stochastically loaded 0.89 pL droplets with ∼0.01 crystals/droplet (∼⍰0.01) [Fig. 1(c)]. Multiple nucleation events within each droplet results in a high crystal size CV. As droplets are miniaturised the mean occupancy falls below ⍰0.1, giving rise to the majority of occupied droplets containing a single crystal. Confirming our expectations, single crystal occupancy promotes uniformity, producing a crystal size CV of ∼15% in 0.89 pL droplets. Importantly, single occupancy coupled with droplet volume control also confers crystal miniaturization, producing ∼3-⍰m-long lysozyme crystals in the smallest, 0.89 pL droplets [Fig. 1(c)].

Attaining single crystal occupancy while aiming to reduce crystal size by limiting droplet volume becomes inefficient as a consequence of the nucleation density. Beyond this, other losses are apparent with droplet miniaturisation [Fig. 1(f)], with the crystal density falling from ∼80 crystals/nL for batch controls and 194 pL droplets to ∼7 crystals/nL in the 0.89 pL droplets. Losses correlate with the increased surface area to volume ratio associated with droplet miniaturization [Fig. 1(f)], which may implicate the surfactant droplet interface as an inhibitory environment for crystal formation. In terms of throughput, losses are compensated by droplet generation frequency increasing with droplet miniaturisation. In practice, droplet generation frequency increases 50-fold from 0.44 kHz with the 754 pL droplets to 23.5 kHz with the 0.89 pL droplets [Fig. 1(g)]. Such throughput, with incubation off-chip, allows the mass production of crystals which is otherwise greatly limited by device size when undertaking on-chip crystallisation.

### 3.3 Pdx1 crystallisation in microfluidic droplets with seeding

We next sought to investigate whether the droplet approach could be applied to a protein with more typical crystallisation behaviour than lysozyme. We used Pdx1, where nucleation rates are much lower, resulting in only a few crystals, inadequate for populating small droplets with crystals. To address this issue, we prepared Pdx1 seeds to substantially increase the crystal density and synchronise crystal growth initiation.

In batch conditions the addition of seeds produced a crystal density of 10^7^/mL with an average length of ∼11 ⍰m [Fig. 2(a)]. Accordingly, ten-fold seed dilution in mother liquor reduced the number of crystals while providing more protein per crystal, resulting in ∼18-⍰m-long crystals for 1/10 seed dilutions and 30-⍰m-long-crystals for 1/100 seed dilutions [Fig. 2(a)]. In principle, seeding initiates crystal growth at the same time, providing equal access to protein throughout growth which results in same-sized crystals. In practice, crystals were variable in size, with a ∼25% CV across the dilution series [Fig. 2(a)].

**Figure 2.**
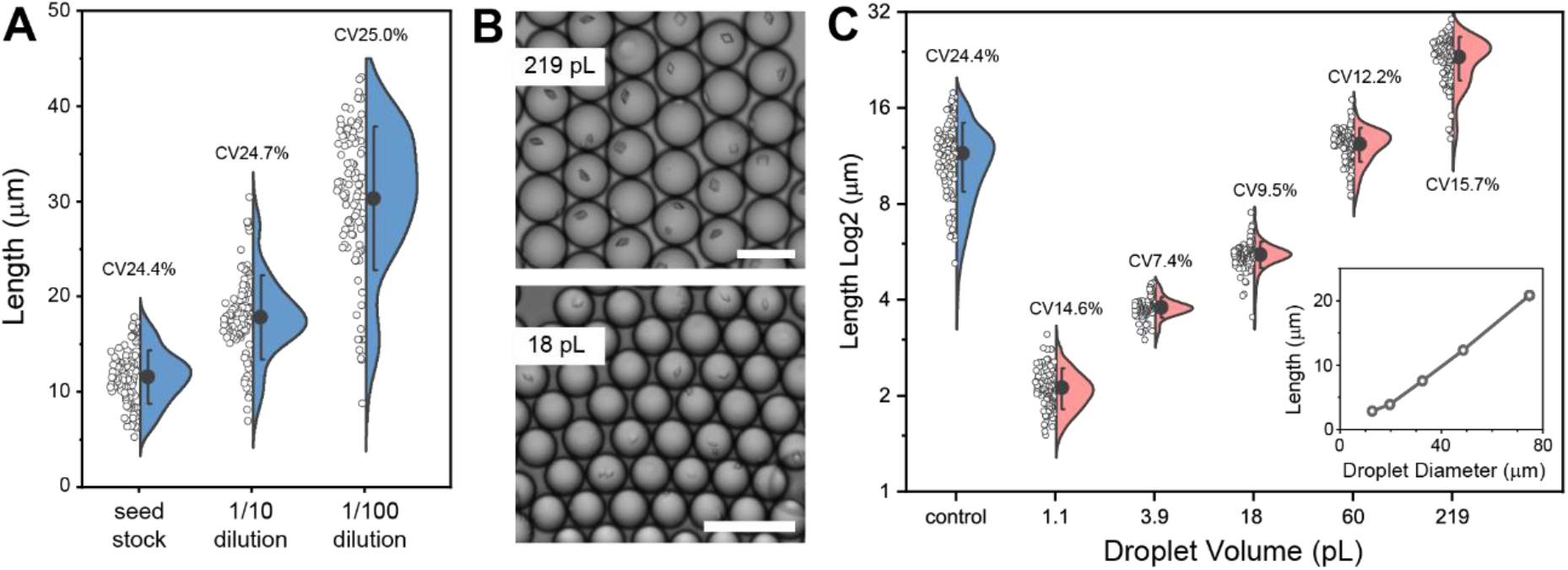
The effect of seeding in batch conditions compared with droplet conditions on Pdx1 crystal size. (a) Seeded batch Pdx1 crystallisation involved a 1:1:1 mixture of Pdx1, seed (10^5^– 10^7^/mL) and mother liquor. The seed dilution effects crystal size (blue), but not crystal uniformity. (b) Pdx1 crystals were grown in droplets using a 2:1 mixture of seeds (10^7^/mL) in mother liquor with Pdx1. Pdx1 crystals grown in 219 and 18 pL monodisperse droplets typically have single occupancy (scale bars 100 ⍰m). (c) Droplet miniaturisation over a 200-fold range was used to control Pdx1 crystal length from ∼20 to ∼2 ⍰m (salmon), with crystal length being proportional to droplet volume. (c, inset) Linear scaling of crystal length with droplet diameter. Droplet confinement enables crystal-size uniformity (CVs 7.4–15.7%). Pdx1 crystals prepared in batch (control, blue) are large with low uniformity (CV 24.4%).

Translating the seeded crystallisation of Pdx1 in batch conditions to droplet environments using 10^7^/mL seeds typically resulted in single crystal occupancy to favour crystal length uniformity (CV 7– 16%) across a 200-fold range of droplet volumes (1.1–219 pL) [Figs. 2(b) and 2(c)]. Droplet volume scaling with single crystal occupancy allows crystal size to be controlled, from ∼2 ⍰m in length for the smallest 1.1 pL droplets, to ∼20 ⍰m in length for the largest 219 pL droplets [Fig. 2(c)]. Overall crystal size can be engineered by droplet volume while retaining uniformity, albeit with crystal occupancy decreasing with diminishing droplet volumes.

### 3.4 Considerations for crystallisation in microfluidic droplets

#### 3.4.1 Aspect ratio

The general applicability of droplets as environments for preparing a variety of different protein crystals is supported by previous work (Heymann *et al*., 2014; Akella *et al*., 2014; Babnigg *et al*., 2022). To extend applicability, we sought to investigate the effect of droplet confinement on the growth of crystal needles. Using trypsin type I as a model needle system, it was evident that droplet diameters are insufficient to allow full elongation, resulting in lower crystal axial ratio (*l*/*w*) or fragmentation into multiple small needle crystals [Fig. S4(a) and S4(b)]. A similar effect is evident with parallelepiped-shaped lysozyme crystals, with crystal axial ratio decreasing with droplet diameter [Fig. S4(c)]. This indicates that protein inclusion within the ends of elongated crystals is impeded within droplets.

#### 3.4.2 Viscosity

Another consideration for the broader utility of droplet microfluidics for crystal preparation is the use of different crystallisation mixtures. Precipitating agents such as poly(ethylene glycol) increase viscosity, which impacts the feasibility of producing stable droplet flows at sufficient throughput. To evaluate this effect, we prepared PEG 6000 solutions (0–25% (w/v)) ranging in viscosity from 1 to 21 mPa·s and used these to observe the effect of viscosity on the generation of 50-m-diameter droplets. Only a ∼3-fold reduction in throughput was observed over these extremes [Fig. S5], indicating scope to apply droplet microfluidics to other crystallisation conditions.

#### 3.4.3 Minimum Crystal Size

The minimum crystal size is another consideration. Given that diffraction data can be obtained from sub-micron crystals (Gati *et al*., 2017, Bücker *et al*., 2020, Williamson *et al*., 2023), and the 2–3 m long lysozyme and Pdx1 crystals prepared in ∼1 pL droplets, there is scope to further reduce droplet volumes. While it is feasible to prepare monodisperse 5.4-m-diameter droplets with a volume of 82 fL, the effect of greatly reduced seed occupancy and lower crystal formation frequency [Fig. 1(f)], prevented observable crystal formation [Fig. S6].

Smaller crystals are also harder to hit with a microfocus X-ray beam and impact sample delivery choice. For instance, small crystals will pass through 7 µm and larger apertures on fixed targets (Hunter *et al*., 2014; Roedig *et al*., 2015; Owen *et al*., 2017; Mehrabi *et al*., 2020), although smaller apertures are now emerging (Carrillo *et al*., 2023). As an alternative, wells within fixed targets can be loaded by depositing 10–100’s pL droplets containing microcrystals using a piezoelectric injector. These droplets are larger than the aperture and held by surface tension to the well walls during data collection (Davy *et al*., 2019).

### 3.5 Serial synchrotron crystallography (SSX)

We tested the visually similar crystals of lysozyme and Pdx1 prepared in batch and droplets, for diffraction power. While droplets can be directly dispensed on silicon fixed targets (Babnigg *et al*., 2022), we opted to remove the fluorinated oil and surfactant to ensure optimal signal to noise. This can be achieved by a procedure called *breaking the emulsion* [see Methods, compare Fig. S2].

SSX experiments were performed at the new ID29 serial beamline at ESRF. Data collection took 10 mins with minimal sample consumption of 3–5 L volumes using the ESRF sheet-on-sheet (SOS) chip sample holder (Doak *et al*., 2018). A full data set was achieved from a single chip of lysozyme with a microcrystal concentration of 10^8^/mL. However, Pdx1 required three chips to obtain complete data, owing to the lower microcrystal concentration of 10^7^/mL and its lower symmetry *H3* space group. Data collection statistics are shown in Table 1.

We used a similar number of integrated lattices to compare data quality, and chose the same resolution cut-off (1.8 Å for lysozyme and 2.5 Å for Pdx1). Data between batch and droplet crystallisation are equivalent for CC_1/2_ and CC^*^ indices, whilst gains in signal-to-noise ratio and R_split_ are observed for crystals grown in droplets [Table 1, Table S4 and Fig. S3].

#### 3.6.1 Mixing in droplets

Substrate-triggered time-resolved experiments require the mixing of crystal and substrate volumes. The median *k*_cat_ for enzymes is 13.7 s^-1^ (∼70 ms reaction cycles) (Bar-Even *et al*., 2011), requiring mixing and into-crystal transport (and binding) times of a few milliseconds to synchronise reactions and allow intermediates to be effectively resolved. However, mixing in conventional microfluidic systems is slow, limited by substrate diffusion into the crystal stream. In contrast, microfluidic droplets lend themselves to fast mixing (Song & Ismagilov, 2003). Here, the transport of droplets in microchannels introduces circulations within the droplet for rapid, convective-diffusive mixing [Fig. S7]. We went on to explore the merits of two different droplet-based mixing approaches.

#### 3.6.2 Mixing by droplet generation and transport

The first system involves mixing by droplet generation and transport. Experiments involved the droplet encapsulation of a stream of pre-formed ∼7×2 ⍰m lysozyme crystals (∼10^7^/mL) with a stream of red dye (SAA, 570 Da), comparable to a typical small molecule substrate. Image analysis reveals that crystal and dye mixing during droplet generation occurs in a stepwise fashion: First laminar streams converge with diffusion between streams initiating slow mixing, then droplet generation causes stream thinning (with short diffusion paths) for rapid mixing, followed by droplet transport with internal circulations driving mixing to completion [Fig. 3(a)]. Mixing begins upon flow convergence, with full mixing defined by a pixel intensity CV of 5%.

**Figure 3.**
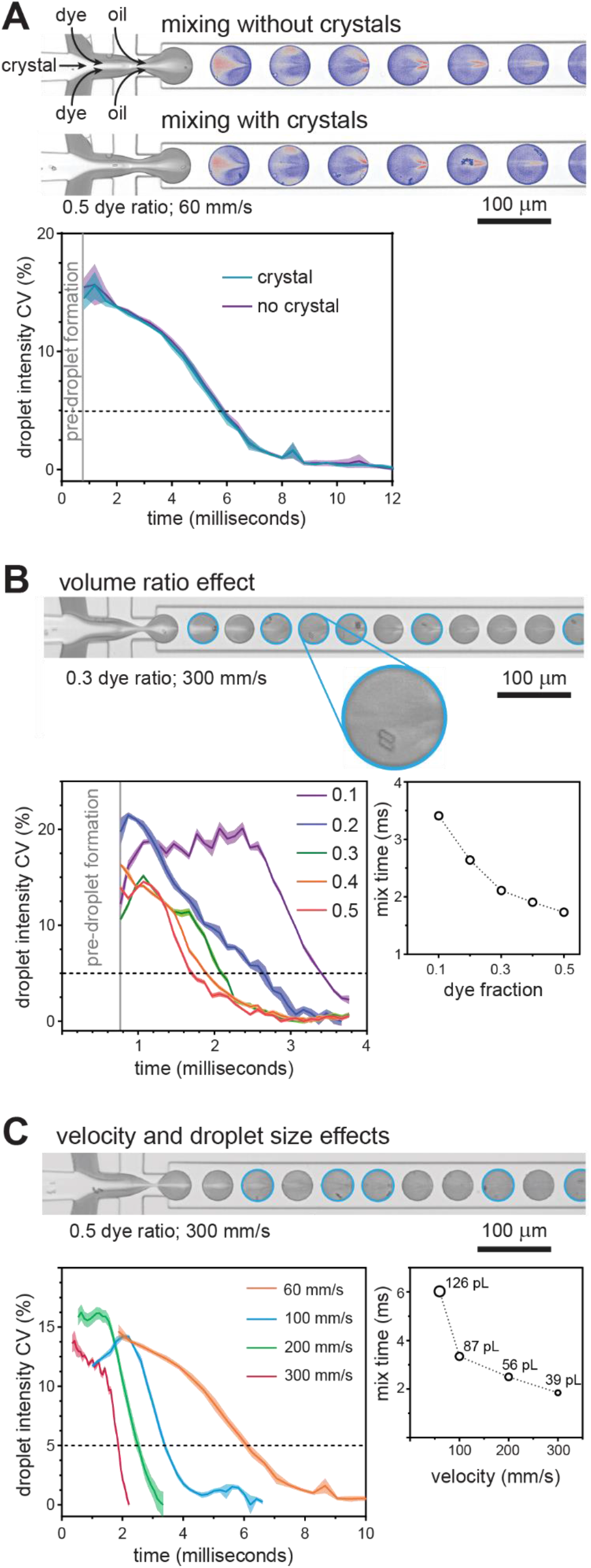
Mixing lysozyme crystals in droplets. (a) Mixing during droplet generation and transport along the channel. The mixing rate with and without crystals is the same (droplets are colour enhanced to aid visualisation). Analysis involved 12 droplets with crystals and 10 droplets without. (b) Dye and crystal mixing in droplets at a ratio of 0.3 with a droplet velocity of 300 mm/s. Droplets containing crystals are highlighted with cyan circles. The mixing rate increases as the volume fraction of dye increases, with the optimal ratio being 0.5 (300 mm/s droplet velocity). The droplet pixel intensity CV is plotted as mean±SD for 15 droplets. (c) Dye and crystal mixing in droplets at the optimal 0.5 ratio with a droplet velocity of 300 mm/s. The droplet pixel intensity CV is plotted as mean±SD for 15 droplets. Increasing velocity increases convection (circulations within droplets) and shrinks droplet volumes to reduce diffusion paths, with both causing faster mixing.

The presence of crystals within droplets did not affect mixing [Fig. 3(a)]. We next tested the ‘entropy of mixing’ theory (Ott & Boerio-Goates, 2000), which states that mixing is maximized when the volumes of initially separate liquids are equal. Indeed, at the same droplet velocity (300 mm/s), mixing times are reduced from 3.4 to 1.73 milliseconds as the volume fraction of dye increases from 0.1 to 0.5 [Fig. 3(b)]. Taking this further we investigated the effect of droplet velocity on mixing using the optimal 1:1 crystal:dye ratio. Increasing the velocity from 60 to 300 mm/s (droplet generation velocity limit) increased circulation speeds within droplets. Higher velocities also impart higher shear stresses during droplet generation, decreasing the droplet volume from 126 to 39 pL and producing shorter diffusion paths. Consequently, mixing times decrease from 6.0 milliseconds at 60 mm/s to 1.85 milliseconds at 300 mm/s [Fig. 3(c)], with faster mixing times anticipated using smaller and higher velocity droplets. While mixing times are seldom reported, such fast mixing is equivalent to high velocity co-axial capillary mixers (Calvey *et al*., 2016), and exceeds mixing by drop-on-drop dispensing (Butryn *et al*., 2021), or the ∼20 millisecond mixing times reported for 3-D printed GDVN devices incorporating mixing blades (Knoška *et al*., 2020).

#### 3.6.2 Mixing initiated by droplet fusion

The ability to produce crystals in droplets affords an alternative strategy for mixing; Protein crystals can be prepared in droplets by incubation (*e*.*g*. overnight) with droplets subsequently injected into a droplet device for fusion with substrate-containing droplets. This removes the need for breaking the emulsion, and moreover droplet-containment prevents crystal sedimentation within the syringe and prevents channels being clogged. As a proof of principle, we developed a microfluidic circuit for generating 225 pL substrate droplets and synchronizing these with pre-formed 200 pL droplets containing crystals (Video S2). Synchronized droplet coupling was achieved by exploiting the size-dependent velocity differences between crystal-containing and substrate-containing droplets: The smaller, faster, droplets approach and contact the larger droplets in readiness for fusion. A surfactant exchange method was used for fusing droplets and initiating mixing (Mazutis *et al*., 2009). Unlike mixing by droplet generation, this does not include the stream thinning effect for shortening diffusion paths. At a droplet velocity of 100 mm/s which enables reliable droplet fusion, this approach achieves mixing in ∼7 milliseconds [Fig. S8, Video S2 and S3], again faster mixing is anticipated for smaller droplets. It is worth noting, that into-crystal substrate transport can occur earlier since convection within the droplet mobilises the crystal throughout substrate-occupied regions before complete mixing is achieved. Nevertheless, the mixing times we report provide a conservative guide for the millisecond timescales that can be accessed, with the limiting step now being the into-crystal travel timescales of the substrate.

#### 3.6.2 Droplets interfacing with the beam

To perform time-resolved experiments, mixing is followed by defined incubations and then crystal interaction with the beam. Importantly, droplets, and crystals within them, have the same transport velocity ensuring uniform incubation times. In contrast, conventional microfluidic transport suffers the effects of the parabolic velocity profile in which crystals in different streamlines are transported at different velocities (*i*.*e*. have different incubations). Periodic droplet generation with tuneable frequency (*e*.*g*. 200 Hz to 6 kHz in our reported mixing experiments) further offers potential for synchronisation with the mean to improve the hit rate. In practice, however, retaining periodicity during ejection into the beam introduces technical challenges which currently limit their potential (Echelmeier *et al*., 2019 and 2020; Doppler *et al*., 2021; Sonker *et al*., 2022). Droplet methods still exceed hit rates achieved using conventional GDVN methods, but now offer the benefits of faster micromixing for synchronised reaction triggering.

Alternatively, the current droplet microfluidic devices offer a means for investigating catalytic processes which have a spectral read-out. This provides a route to experiment work-up in advance of visiting synchrotron or XFEL facilities. To exploit synchrotron capabilities and have broad utility new challenges and technical possibilities emerge such as the fabrication of droplet microfluidic devices using thin-film materials (*e*.*g*. cyclic olefin co-polymer) that do not appreciably attenuate the X-ray beam (Sui *et al*., 2016; Liu *et* al., 2023). For much higher energy XFEL sources the challenge of controlled ejection into the beam remains.

## 4. Conclusion

In this paper we have demonstrated droplet confinement and miniaturization for controlling crystal size and uniformity. At low picolitre and femtolitre scales nucleation becomes improbable and can be bypassed using a seeding strategy for producing crystals only a few microns in length. The method was demonstrated with lysozyme and Pdx1, with crystals grown in droplets producing equivalent quality diffraction data to those produced in batch conditions. Picolitre-scale droplet microfluidics also enables rapid, millisecond-scale micromixing to increase the temporal resolution of time-resolved experiments. Droplet microfluidic mixers can, in the future, be fabricated using thin-film, X-ray transparent materials for synchrotron experiments or coupled with beam injection methods to extend the approach to XFEL experiments. In summary, droplet microfluidics methods offer great promise for improving time-resolved crystallography.

## Acknowledgements

We thank Chris Holes for support with protein crystallisation, ESRF for access to ID29 (under proposal numbers MX2438/MX2551) and Emma Beale for the adapted lysozyme crystallisation conditions.

## Funding Information

The research was funded by a Diamond Doctoral Studentship Programme (JS, RB), a South Coast Biosciences Doctoral Training Partnership SoCoBio DTP BBSRC Studentship BB/T008768/1 (JS), Wellcome Investigator Award 210734/Z/18/Z (AMO), Royal Society Wolfson Fellowship RSWF\R2\182017 (AMO), EPSRC Impact Acceleration Account and EPSRC Transformative Healthcare grant EP/T020997/1 (NH), University of Southampton Seed Enterprise Development Fund (NH), BBSRC BB/S008470/1 (IT) and STFC “Serial Data Processing and Analysis in CCP4i2 (IT, MM).

## Notes

### Competing Interest Statement

The authors have declared no competing interest.

