## Supplementary Information for "Droplet microfluidics for time-resolved serial crystallography"

### Supporting information

**Table S1** Droplet microfluidic conditions for preparing lysozyme crystals.

| Junction Dimensions (*w, h*) (see Fig. 1A) | QX200 Fluoro-oil | Mother Liquor | Lysozyme | Droplet Diameter | Droplet Volume | Droplet Frequency |
|---|---|---|---|---|---|---|
| 125x100 µm | 60 µL/min | 16 µL/min | 4 µL/min | 112.9 µm | 754 pL | 0.44 kHz |
| 70x75 µm | 37.5 µL/min | 10 µL/min | 2.5 µL/min | 71.8 µm | 194 pL | 1.07 kHz |
| 50x50 µm | 22.5 µL/min | 6 µL/min | 1.5 µL/min | 54.4 µm | 84.3 pL | 1.48 kHz |
| 35x35 µm | 15 µL/min | 3 µL/min | 0.75 µL/min | 33.5 µm | 19.7 pL | 3.18 kHz |
| 22x20 µm | 8 µL/min | 2 µL/min | 0.5 µL/min | 23.2 µm | 6.54 pL | 6.37 kHz |
| 20x12 µm | 8 µL/min | 1.6 µL/min | 0.4 µL/min | 17.6 µm | 2.85 pL | 11.7 kHz |
| 10x12 µm | 5 µL/min | 1 µL/min | 0.25 µL/min | 11.9 µm | 0.89 pL | 23.3 kHz |

**Table S2** Droplet microfluidic conditions for preparing Pdx1 crystals.

| Junction Dimensions (*w, h*) | QX200 Fluoro-oil | Mother Liquor + Seed | Pdx1 | Droplet Diameter | Droplet Volume | Droplet Frequency |
|---|---|---|---|---|---|---|
| 70 x 75 µm | 90 µL/min | 26.67 µL/min | 13.33 µL/min | 74.8 µm | 219 pL | 3.04 kHz |
| 50x50 µm | 60 µL/min | 13.33 µL/min | 6.67 µL/min | 48.6 µm | 60.1 pL | 5.54 kHz |
| 35x35 µm | 40 µL/min | 6.67 µL/min | 3.33 µL/min | 32.5 µm | 18.0 pL | 9.27 kHz |
| 20x12 µm | 16 µL/min | 2.67 µL/min | 1.33 µL/min | 19.6 µm | 3.94 pL | 16.9 kHz |
| 10x12 µm | 8 µL/min | 1.33 µL/min | 0.67 µL/min | 12.7 µm | 1.07 pL | 31.1 kHz |
| 10x5 µm | 2x1 µL/min | 0.2 µL/min | 0.1 µL/min | 5.38 µm | 82 fL | 61.3 kHz |

**Table S3** Droplet microfluidic conditions for preparing trypsin type I needle crystals.

| Junction Dimensions (*w, h*) | QX200 Fluoro-oil | Mother Liquor | Trypsin | Seeds | Droplet Diameter | Droplet Volume | Droplet Frequency |
|---|---|---|---|---|---|---|---|
| 70 µm | 54 µL/min | 6 µL/min | 6 µL/min | 6 µL/min | 75 µm | 221 pL | 1.36 kHz |
| 50 µm | 27 µL/min | 3 µL/min | 3 µL/min | 3 µL/min | 50 µm | 65 pL | 2.29 kHz |
| 35 µm | 9 µL/min | 1 µL/min | 1 µL/min | 1 µL/min | 35 µm | 22 pL | 2.22 kHz |

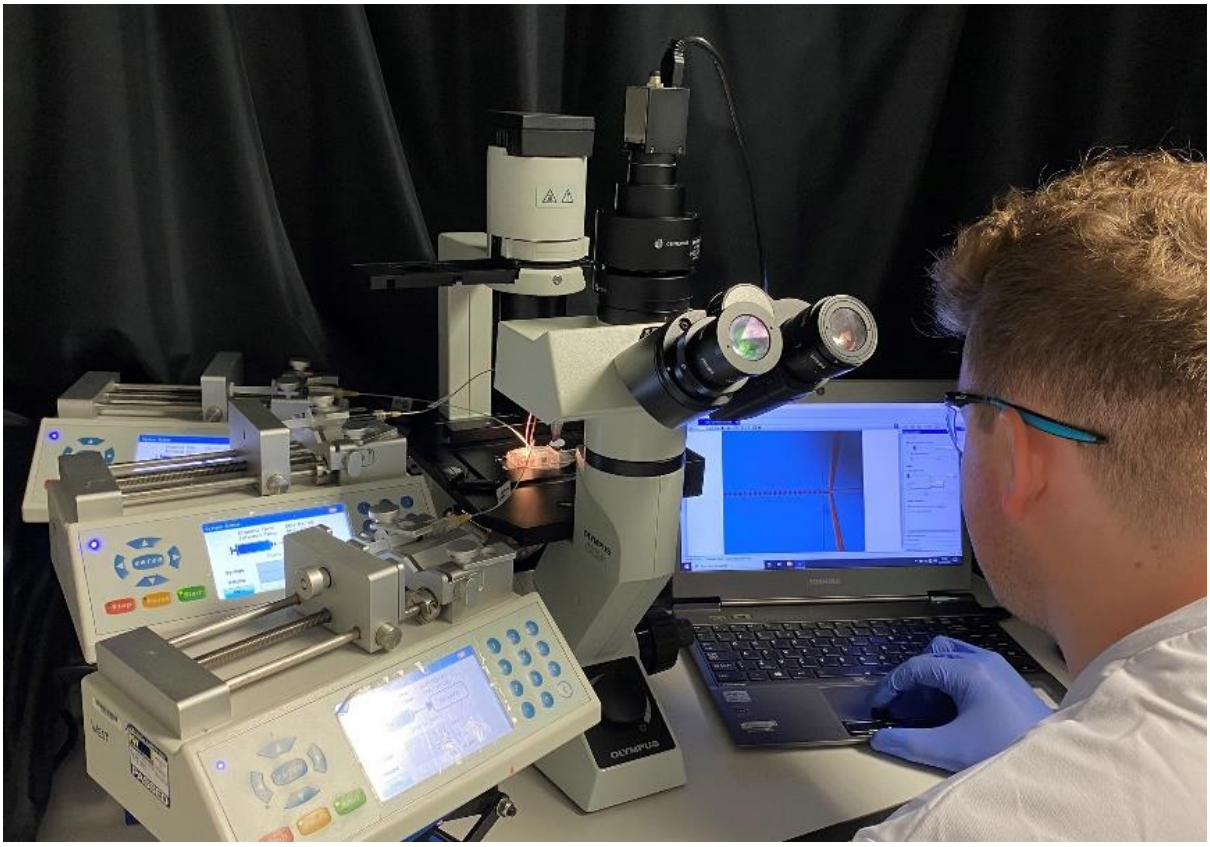


**Figure S1** Droplet microfluidics experimental setup.

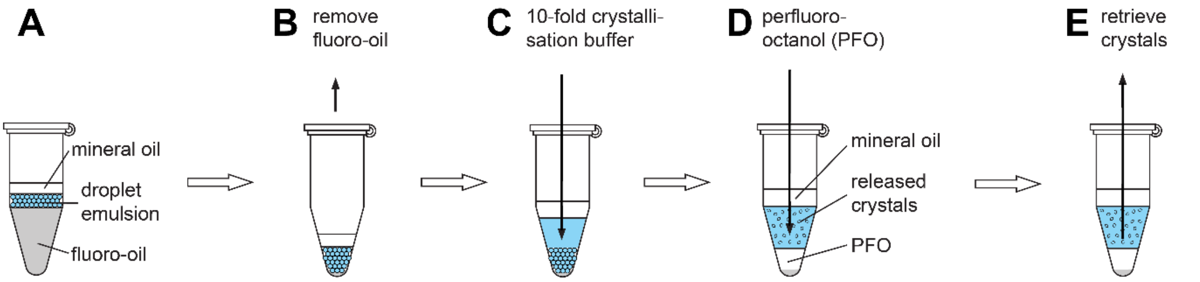


**Figure S2** Breaking the emulsion. Droplets are collected under a mineral oil overlay, with the buoyant nature causing an emulsion to form above the fluoro-oil carrier (A). The fluoro-oil is removed (B), and a ~10-fold volume of crystallisation buffer relative to emulsion volume is added to the emulsion (C). Next, a 1-fold volume of perfluoro-1-octanol (PFO) is added, and gently mixed to coalesce all droplets into a single aqueous volume (D). This volume and the crystals within it are retrieved for analysis (E).

**Serial Synchrotron Crystallography (SSX) Data Collection and Processing**

A hit rate of 41.6% was achieved for the lysozyme control preparation and 27.9% for the droplet preparation, compared to a hit rate of 8.2% for the Pdx1 control preparation and 8.5% for the droplet preparation. Detector geometry refinement was performed with *geoptimiser* (Yefanov *et al.,* 2015) and a smaller subset of 1000 images for each dataset, ensuring high indexing rates. Lysozyme control preparations produced 29,954 indexable patterns, resulting in 5,990 indexed patterns/µL, and droplet preparations produced 22,304 indexable patterns, resulting in 4,460 indexed patterns/µL. In comparison, Pdx1 control preparations produced 19,325 indexable patterns, resulting in 1,288 indexed patterns/µL, and droplet preparations produced 20,635 indexable patterns respectively, resulting in 1,375 indexed patterns/µL. The number of integrated lattices is much greater than those indexed, especially in the case of lysozyme where multiple lattices were observed in the same shot. During data processing, the effect of multiple lattices was tested by including or omitting the --multi argument in the *indexamijig* command. The inclusion of multiple lattices had no detrimental effect on data quality.

Diffraction patterns were processed using CrystFEL (v.0.10.2; White *et al.,* 2012, 2019). Images were stored in a hdf5 stream and initial hit finding was performed with Cheetah (Barty *et al.,* 2014). Hits were indexed with x*gandalf* (Gevorkov *et al.,* 2019) and integrated using the following parameters: --peaks=peakfinder8 --multi --int-radius=4.0,6.0,10.0 --tolerance=5,5,5,1.5,1.5,1.5 --peak-radius=4,6,10 --min-peaks=30 --min-snr=4.0 --threshold=1500 --local-bg-radius=5 --min-res=80 --max-res=1200 --min-pix-count=3 --max-pix-count=200. Indexing ambiguities present in the Pdx1 datasets (arising from the *H*3 space group) were resolved using *ambigator* and the following parameters: -y 3_H --operator=k,h,-l --iterations=20 --highres=3.0. Data were merged using *partialator* with partialities, post-refinement and scaling using the following parameters: -y 4/mmm (Lysozyme) or 3_H (Pdx1) --model=xsphere --iterations=1. Figures of merit and .mtz files for crystallographic structure determination were generated using the import_serial task, currently available in the latest version of CCP4 (8.0.016) (Agirre *et al.,* 2023) (https://github.com/MartinMalyMM/import_serial).

**Table S4** Data collection statistics for lysozyme and Pdx1 crystals grown in batch and in droplets. Data collected at ESRF/ID29. Values in parentheses are for the outermost resolution shell.

| | **Lysozyme Control** | **Lysozyme Droplet** | **Pdx1 Control** | **Pdx1 Droplet** |
|---|---|---|---|---|
| Average crystal length | ~15 mm | ~15 mm | ~15 mm | ~15 mm |
| No. of collected images | 81,800 | 81,800 | 245,400 | 245,400 |
| No. of hits | 34,032 | 22,815 | 20,268 | 20,827 |
| Hit rate (%) | 41.6 | 27.9 | 8.2 | 8.5 |
| Indexed images (single lattice) | 29,954 | 22,304 | 19,325 | 20,635 |
| Indexing rate (%) | 88.0 | 97.7 | 95.3 | 99.0 |
| Integrated patterns (including multiple lattices) | 58,984 | 51,812 | 27,581 | 25,464 |
| Space group | $P4_32_12$ | $P4_32_12$ | $H3$ | $H3$ |
| $a = b$ (Å) | 79.0 | 78.9 | 177.9 | 180.3 |
| $c$ (Å) | 37.9 | 37.9 | 117.3 | 119.2 |
| α, β, γ (˚) | 90, 90, 90 | 90, 90, 90 | 90, 90, 120 | 90, 90, 120 |
| Resolution (Å) | 79.00 – 1.80 (1.83 – 1.80) | 78.90 – 1.80 (1.83 – 1.80) | 93.33 – 2.50 (2.54 – 2.50) | 94.75 – 2.50 (2.54 – 2.50) |
| Total reflections | 5,096,019 (26,983) | 5,385,630 (28,772) | 4,534,670 (215,526) | 4,424,291 (210,986) |
| Unique reflections | 11,636 (669) | 11,607 (663) | 47,878 (4703) | 50,011 (4635) |
| Completeness (%) | 100.0 (100.0) | 100.0 (100.0) | 100.0 (100.0) | 100.0 (100.0) |
| Multiplicity | 438.0 (47.67) | 464.0 (51.94) | 47.4 (45.18) | 44.2 (41.83) |
| $\langle I/\sigma(I) \rangle$ | 13.5 (0.2) | 17.2 (0.9) | 4.4 (0.5) | 5.0 (0.8) |
| $CC_{1/2}$ | 0.99 (0.49) | 0.99 (0.44) | 0.96 (0.17) | 0.96 (0.27) |
| $CC^*$ | 0.99 (0.81) | 0.99 (0.78) | 0.99 (0.54) | 0.99 (0.65) |
| $R_{split}$ | 5.2 (343.7) | 4.9 (100.8) | 19.6 (212.2) | 17.1 (147.3) |

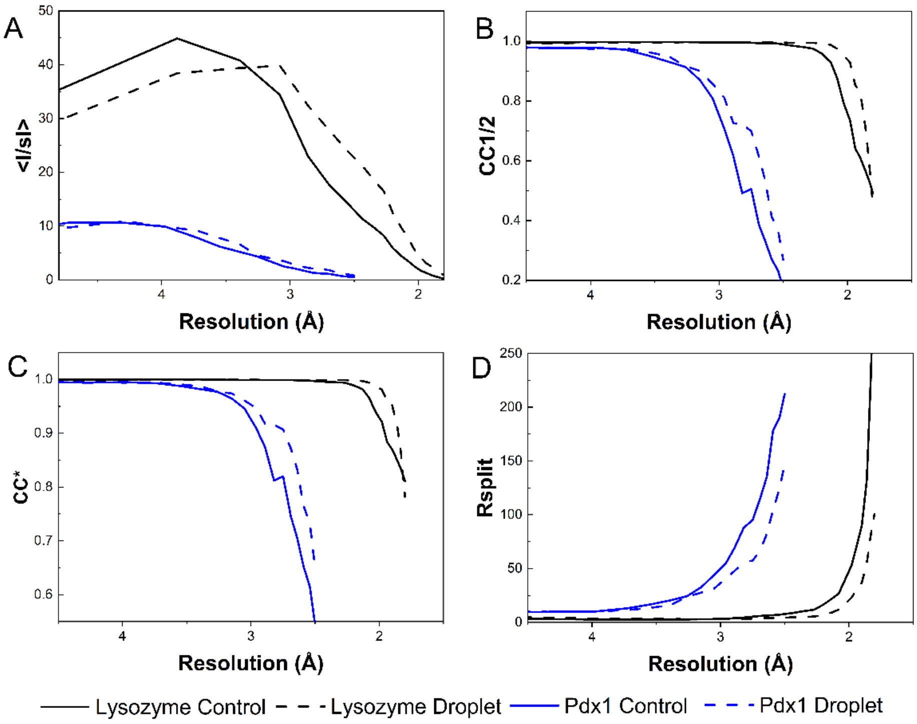


**Figure S3** Plots of figures of merit. (A) Mean signal-to-noise ratio I/sigma(I), (B) Pearson's correlation coefficient CC1/2 , (C) CC* and (D) Rsplit against resolution comparing lysozyme and Pdx1 crystals grown in batch with those grown in droplets.

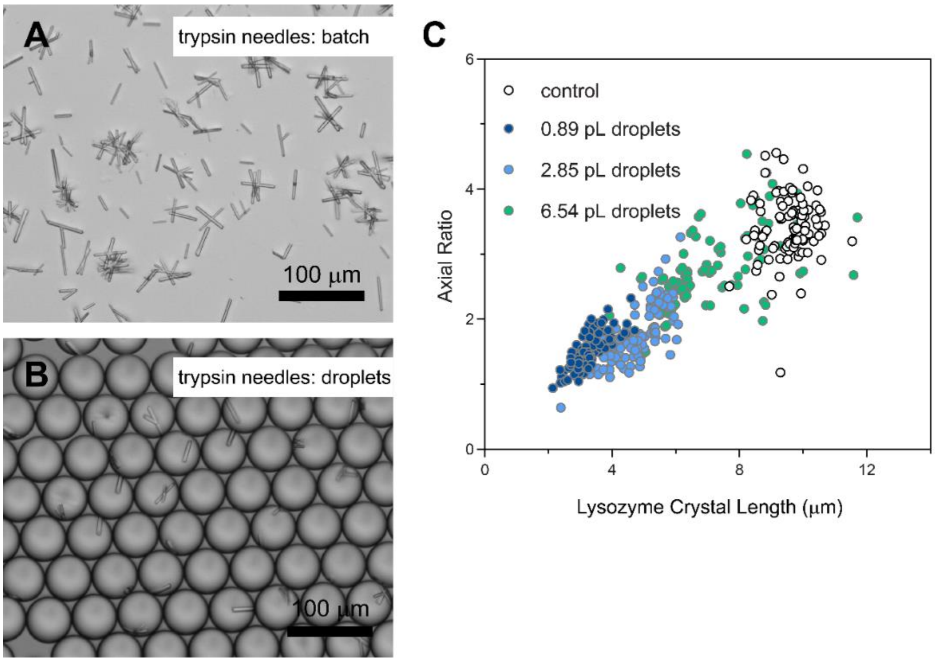


**Figure S4** Trypsin needles and confinement effects on lysozyme axial ratio. (A) Trypsin needles prepared in batch have various lengths with an axial ratio of ~10, (B) whereas crystallisation in droplets produces an axial ratio of ~5. (C) The axial ratio of lysozyme crystals grown in droplets tends to unity as droplets are miniaturised below 1 pL.

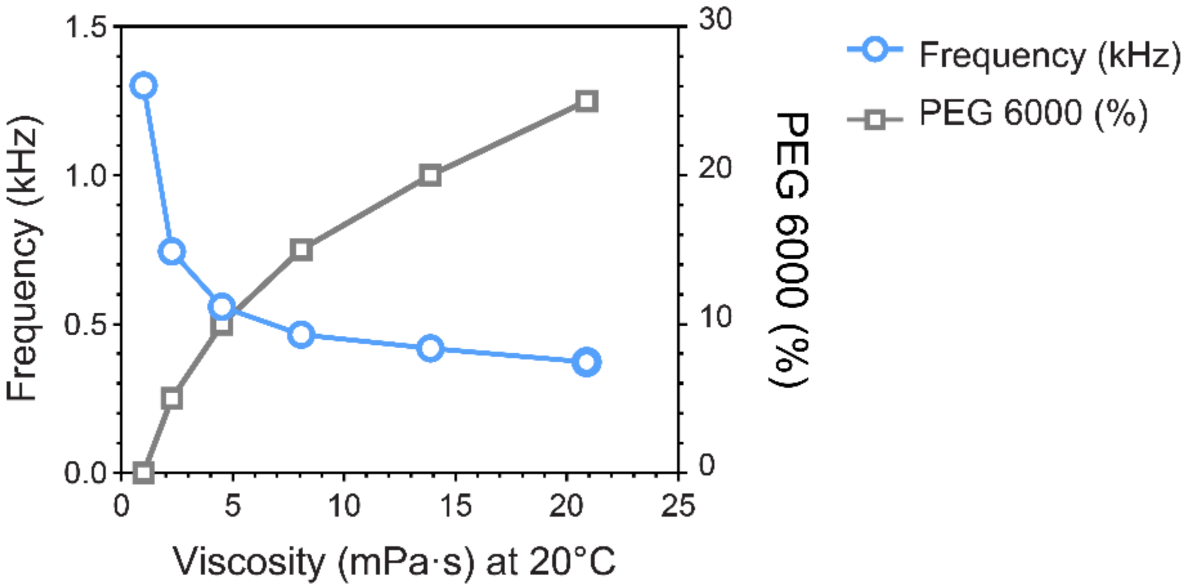


**Figure S5** Viscosity effect on droplet generation throughput. Droplet microfluidics can manage high viscosity crystallisation buffers, although increased viscosity decreases the droplet generation frequency.

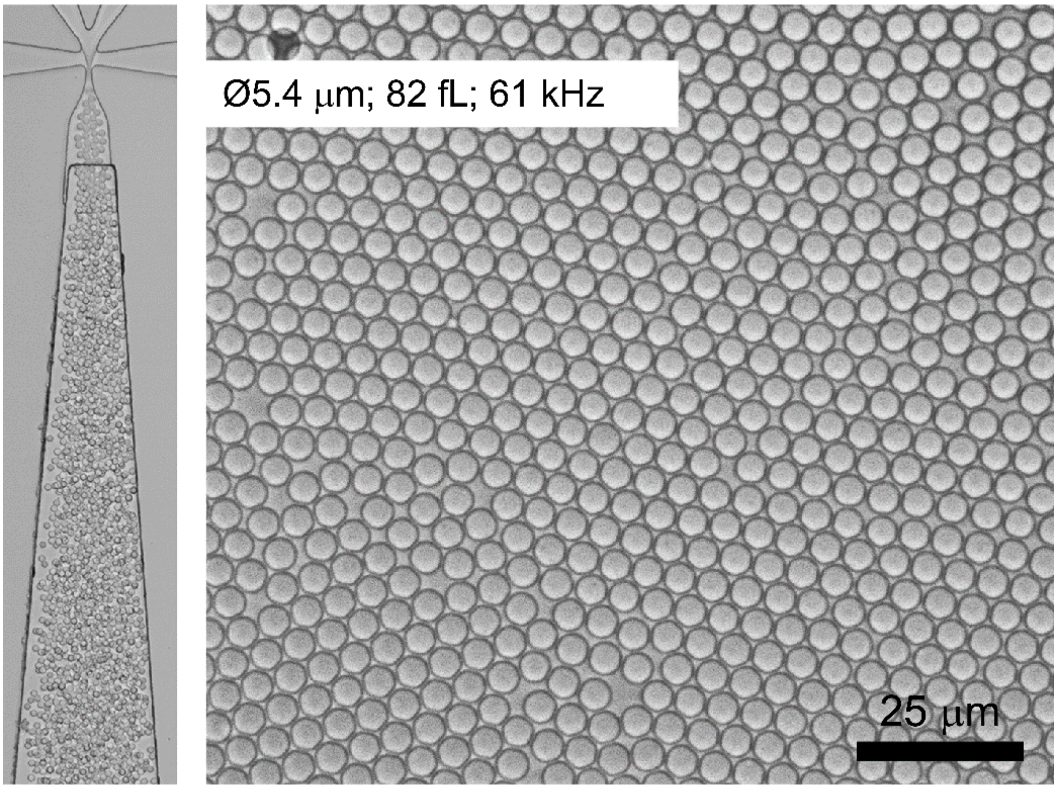

**Figure S6** Femtolitre droplets for Pdx1 crystallisation. Miniaturisation enables the high throughput (61 kHz) generation of ~5-µm-diameter, 82 fL droplets (left). Even with seeding, Pdx1 crystals were not apparent with a 60x/1.4NA oil immersion objective (right).

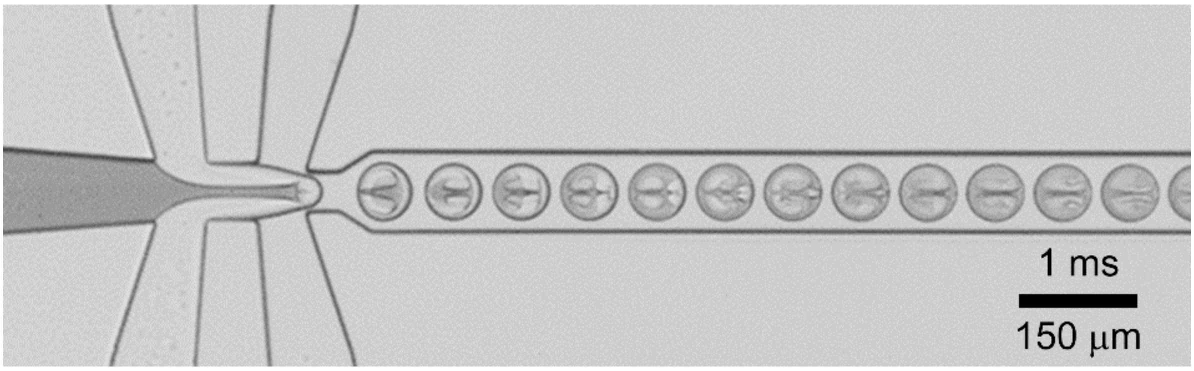


**Figure S7** Circulations within droplets provide convection to drive rapid micromixing.

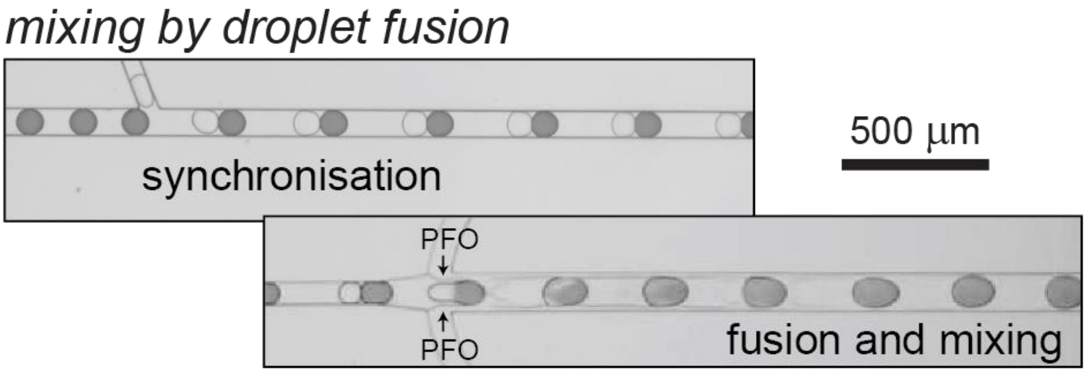


**Figure S8** Droplet synchronisation and 1:1 coupling followed by surfactant exchange using PFO to trigger droplet fusion and mixing (Video S2 and S3). The synchronisation channel is 75x40 mm (*w*,*h*) with a droplet velocity of 120 mm/s. The droplet fuse and mix channel is *w* 125x40 mm (*w*,*h*) with a droplet velocity of 85 mm/s. 200 pL crystal droplets are fused with 225 pL dye droplets with mixing achieved in ~7 milliseconds.
